## Supplementary Material for "Structural comparison of protein-RNA homologous interfaces reveals widespread overall conservation contrasted with versatility in polar contacts"

### Supplementary methods

##### Identification of ribosomal protein-RNA interfaces

As ribosomal complexes are predominant among protein-RNA complexes in the PDB, we wanted to control ribosome bias in our dataset. We used a list of 400 ribosomal protein identifiers (240 human, including human mitochondrial ribosome proteins, 36 from *Escherichia coli*, and 124 from other species, including *Trypanosoma brucei* and *Saccharomyces cerevisiae*) that cover a large part of our ribosomal dataset, as assessed by a simple regular expression search. We used HHblits and HHsearch [1] version 3.0.0 (15-03-2015) to generate an HMM profile for each of these proteins and search for homologous chains in the pdb70 database [1], which we expanded using the redundancy information (pdb70_clu.tsv file) to cover the entire PDB.

Applying this protocol, 615 of our 977 representative protein-RNA interfaces (63%) were assigned as ribosomal and the remaining 362 as non-ribosomal.

##### ECOD classification

We assigned ECOD domains at the T-group level (demonstrating homology and similar topological connections) to all interfaces in our dataset. At the interolog group level, we listed and counted all common ECOD domains of interfaces within each group and attributed a most or a set of most represented ECOD T-group domain(s) to this group using the following criteria. If from all ECOD domain labels within a family, there is a single most represented ECOD domain, which is represented in more than 50% of all family members, then the family is given this domain as a label. If several ECOD domains are equally most represented, and they are represented in strictly more than 50% of all family members, then the family is given these domains as a label. If no ECOD domain label is represented in more than 50% of all family members, this family is labeled as “ambiguous”. Finally, if there are no ECOD domain labels within a family, then the family is unlabeled.

When comparing pairs of interologs, we look at the intersection between labeled ECOD domains of each interface. In the web interface for database exploration, we assign ECOD labels to groups and pairs of interologs.

##### Alternative methods of contact conservation calculation

We tested an alternative generalized Jaccard index [2, 3]. In this setting, contact conservation was determined for a given pair of interologs by calculating the ratio of the sum of the minimum number of atomic contacts over all aligned amino acid-nucleotide pairs divided by the sum of the maximum number of atomic contacts. The contact conservation distribution was then assessed for each pair of interologs in the dataset.

We also computed a non-weighted version of conservation where the calculation does not involve weighting based on contact numbers. Instead, it focuses solely on the presence or absence of contact (defined by heavy-atom minimum distance of 5 Å) in each interface residue-nucleotide pair and its structural equivalent for each pair of interologs.

### Supplementary results

##### Interface analysis and composition

Resulting from the 977 all against all structural alignment, we obtained 2,022 interologs from 765 interfaces. We analyzed this dataset of 765 interfaces after checking that no major differences were observed compared to the dataset of 977 interfaces. Among the 765 interfaces, 33% of the interacting amino acids are in a helix, 10% in a β-strand, and 57% in a "coil" region. Regarding nucleic acid secondary structure, 75% of nucleotides were paired, and 25% were unpaired.

We also assessed the distribution of interface contacts according to the assignment of core/rim protein interface regions: in 55% of the contacts, the amino acid belongs to the core region and in 45% to the rim region.

We analyzed the global number of atomic contacts for each interface. We found that, on average, an interface contains 1160 atomic contacts (vs. 230 for protein-protein interfaces) (1235 for ribosomal interfaces, 987 for non-ribosomal interfaces) and 257 apolar contacts (262 for ribosomal interfaces, 243 for non-ribosomal interfaces). When grouped into residue-nucleotide contacts, an interface has an average of 109 distance-based contacts (120 for ribosomal interfaces, 89 for non-ribosomal interfaces), much more than for protein-protein interfaces, on which the average was 61 [4]. Each interface has on average 2 π-stacking interactions (2 for ribosomal interfaces, 3 for non-ribosomal interfaces) and 30 hydrogen bonds (33 for ribosomal interfaces, 23 for non-ribosomal interfaces). The higher number of stacking interactions in the non-ribosomal interfaces may be explained by different residue type compositions, particularly aromatic amino acids representing 11% of the amino acids in the non-ribosomal interfaces vs. 7% in ribosomal interfaces. We verified that the composition of the interface is consistent with previous statistical analyses performed on smaller sets of heteromeric protein-RNA complexes [5].

Previous studies highlighted that the backbone phosphates are involved in the majority of protein-RNA interactions [6]. It is also the case in our contact dataset, where 59% of the hydrogen bonds involve the phosphate group of the nucleotide. 76% of the amino acids performing an H-bond involve their side chains.

**Supplementary interologs analysis**

Interologs were split into ribosomal and non-ribosomal: interologs were assigned as ribosomal if both protein chains in the structural interolog pairs were assigned as ribosomal protein chains and non-ribosomal when the two protein chains were assigned as non-ribosomal. We decided not to include interologs where one interface was assigned as ribosomal and the other as non-ribosomal; these pairs represented only 2% of our interologs pairs.

Applying these criteria led to the identification of 2,022 interologs, out of which 1371 were ribosomal and 671 non-ribosomal. Those interologs come from 765 interfaces, and those interfaces are made from 257 PDB complexes. 176 (69%) have resolution higher than 2Å.

Among the 2,022 interolog pairs 48 have both interface resolutions lower than 2Å, and in 463 cases, one structural interolog within the pair has resolution lower than 2Å and the other higher than 2Å. In the remaining 1391 cases, both have resolutions higher than 2Å.

The numbers of contacts differ by 19% on average between two interologs for atomic contacts (vs. 15% for protein-protein interfaces) and by 27% for apolar contacts.

Interologs are separated into four equally populated groups of interface sequence identity: the 0-19%, 19-34%, and 34-60% sequence identity groups each contain 505 pairs of interologs, while the 60-100% sequence identity group contains 507 pairs of interologs.

When counting the number of structurally aligned amino acid-nucleotide pairs, where the pair is at a distance shorter than 5 Å in at least one of the two interologs, the 0-19% group has 43342 such pairs, while the 19-34%, 34-60%, and 60-100% have respectively 69337, 62435 and 61786. Hence, the contact conservation in the 0-19% group might be moderately over-estimated because many amino acid-nucleotide pairs cannot be structurally aligned.

**Supplementary contact conservation analysis**

With the alternative generalized Jaccard index, apolar contacts appear slightly less conserved than atomic contacts (S2 Fig). This can be explained by the observation that within a pair of aligned amino acid-nucleotide contacts, the numbers of atomic contacts are more similar than the numbers of apolar contacts (difference of 20% vs. 28% on average); therefore the minimum and maximum numbers of atomic contacts are more similar than the minimum and maximum numbers of apolar contacts and the min/max ratio will be closer to 1 for atomic contacts.

The contact conservation is, on average, 22% when the nucleotide changes base-pairing status between one interolog and the other. To quantify this observation, we analyzed the subgroup of contacts where the nucleotide is base-paired in one interolog and unpaired in the other, constituting 30% of the pairwise contacts. Among this subgroup's 62,182 non-conserved pairwise contacts, 62,032 (99%) are attributed to contacts involving base-paired nucleotides that were lost when the nucleotides transitioned to an unpaired state.

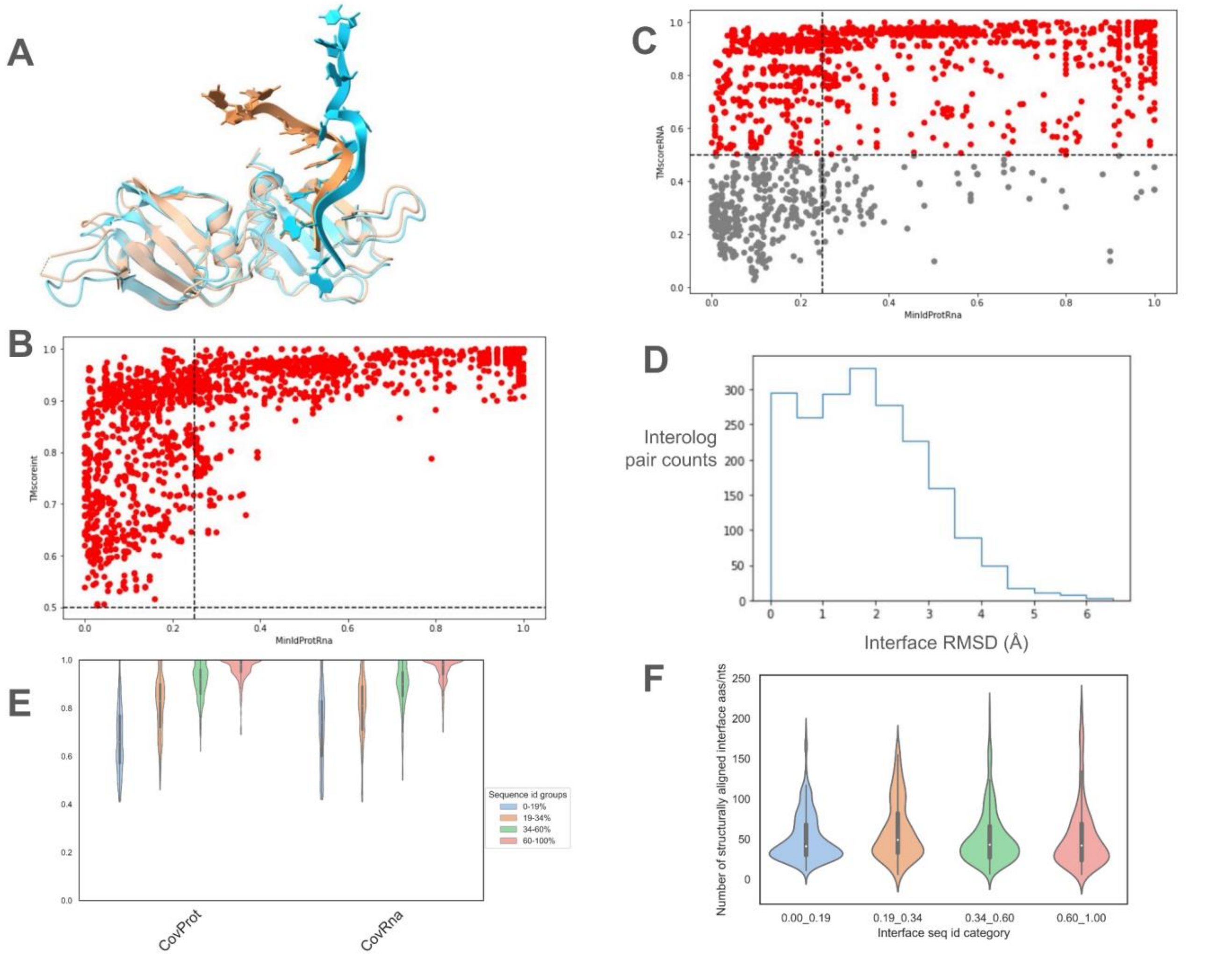


**S1 Fig**: Supplementary information about the construction of the dataset of interologs. **(A)** Structural alignment of 5HO4_A_B and 5WWE_A_B. For this pair, the protein TM-score is 0.96, the RNA TMscore is 0.13, and the interface TMscore is 0.98. Despite low RNA TM-score (due to the flexibility of the RNA molecule), these interfaces are structural interologs. **(B)** Scatter plot depicting the distribution of interface TM-scores depending on the minimum interface sequence identity for the final set of 2,022 interologs. **(C)** Scatter plot depicting the distribution of RNA TM-scores depending on the minimum interface sequence identity for 2,022 final pairs of confidently assigned structural interologs. In gray, 515 pairs have RNA TM-scores below 0.5. This highlights the challenge of using RNA TM-score to define interologs. **(D)** Histogram of interface RMSD (Å) for the 2,022 pairs of interologs. 99.9% have interface RMSD below 6Å and 95.5% have interface RMSD below 4Å. **(E)** Violin plots of interface overlap on protein (CovProt) and RNA (CovRna) for each of the 2,022 interolog pairs, separated into four groups of interface sequence identity (blue: 0-19%, orange: 19-34%, green: 34-60%, red: 60-100%). **(F)** Violin plots of interface intersect size, measured as the number of structurally aligned interface amino acids/nucleotides for each of the 2,022 interolog pairs, separated into four groups of interface sequence identity (blue: 0-19%, orange: 19-34%, green: 34-60%, red: 60-100%).

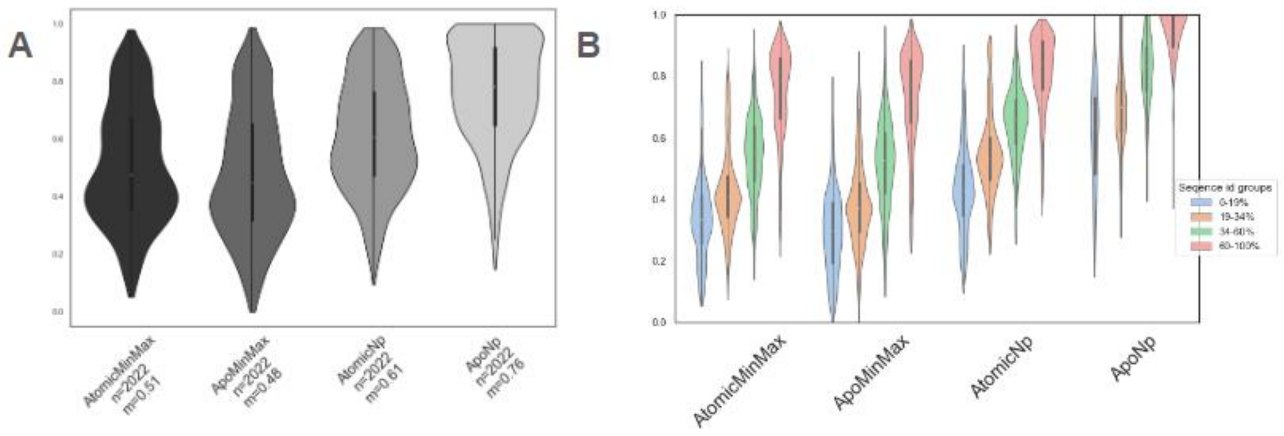


**S2 Fig**: Distributions of contact conservation using alternative metrics. (A) Overall distributions of AtomicMinMax/ApolarMinMax contact conservation (atomic/apolar conservation calculated with generalized Jaccard index using the ratio of smaller to larger number of atomic contacts in each pair of aligned amino acid-nucleotide contacts), and AtomicNp/ApoNp (non-weighted atomic/apolar contact conservation). The difference between AtomicMinMax and ApolarMinMax conservations is statistically significant (p-value = 9.9e-7 in a Wilcoxon rank sum test). The difference between non-weighted atomic/apolar contact conservation is statistically significant (p-value = 4e-129 in a Wilcoxon rank sum test). (B) Distributions of the same contact conservation values as in panel A, separated into four groups of interface sequence identity (blue: 0-19%, orange: 19-34%, green: 34-60%, red: 60-100%). The same trends are observed as in main Figure 3D.

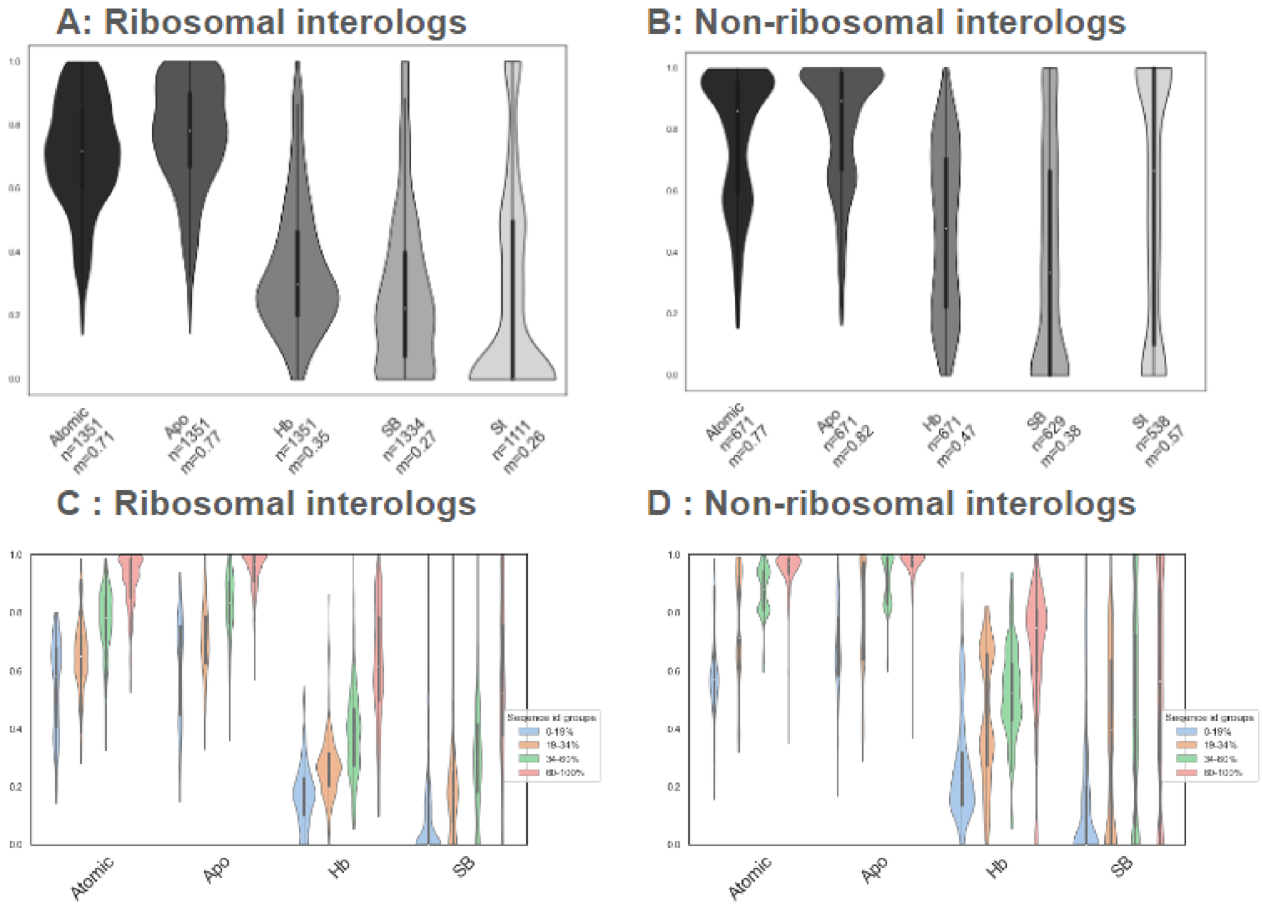


**S3 Fig**: Contact conservation analysis for ribosomal and non-ribosomal interologs. **A & B** Violin plot distribution of contact conservation for atomic contacts (Atomic), apolar contacts (Apo), H bonds (Hb), Stacking contacts (St) for ribosomal interologs (panel A) and non-ribosomal interologs (panel B). n is the number of interolog pairs used in each violin xplot and m is the mean conservation ratio. The p-value between distributions of atomic contacts and apolar contacts denoted by *** in this figure is <1.25e-09 in a Wilcoxon rank sum test. **C & D**: Violin plot distribution of contact conservation for atomic contacts (Atomic), apolar contacts (Apo), H bonds (Hb) for pairs of interologs separated into four groups of interface sequence identity (blue: 0-19%, orange: 19-34%, green: 34-60%, red: 60-100%). For all types of contacts, the differences between any two distributions of conservation among the four groups of sequence identities are statistically significant (p-value<1.6e-19 in Wilcoxon rank sum tests).

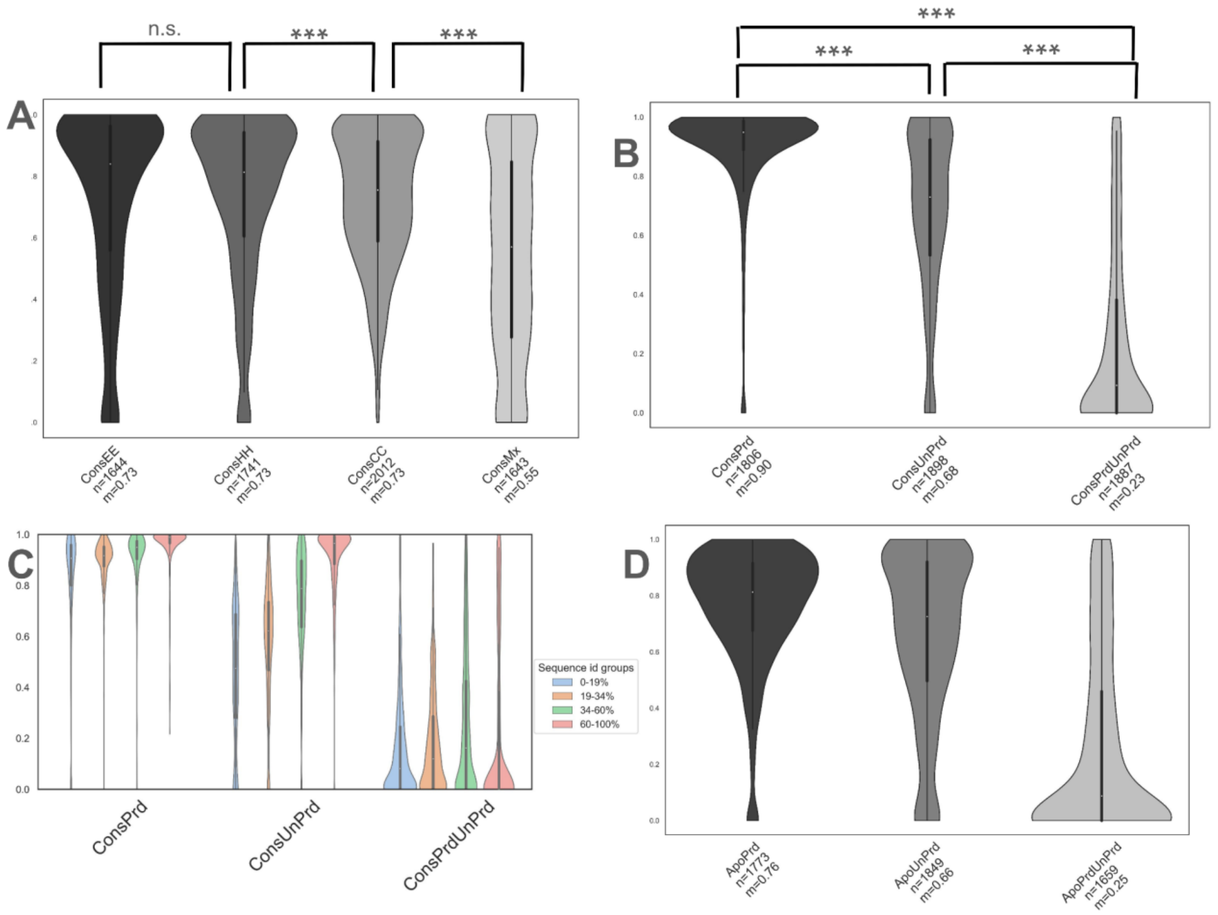


**S4 Fig:** Violin plots of atomic contact conservation depending on residue secondary structure properties. n is the number of interolog pairs used in each violin plot and m is the mean conservation ratio. **(A)** From left to right with a descending gray palette, distributions of atomic contact conservation where the amino-acid and its structural equivalent keep the same type of secondary structure (strand for ConsEE, coil for ConsCC and helix for ConsHH) or have different secondary structure (ConsMx). P-values between distributions denoted by *** in this panel are <2e-05 in a Wilcoxon rank sum test (and n.s. is not significant). **(B)** From left to right with a descending gray palette, distributions of atomic contact conservation where the nucleotide and its structural equivalent have the same secondary structure (base-paired for ConsPrd and unpaired for ConsUnPrd) or change base-pairing status (ConsPrdUnPrd). P-values between distributions denoted by *** in this panel are <2e-166 in a Wilcoxon rank sum test. **(C)** Distribution of apolar contact conservation according to the same categories as in panel C when splitting the interfaces into four groups of interface sequence identity (blue: 0-19%, orange: 19-34%, green: 34-60%, red: 60-100%). **(D)** Distribution of apolar contact conservation in the subgroup where both structurally aligned nucleotides are base-paired (ApoPrd), where both structurally aligned nucleotides are unpaired (ApoUnPrd) or where one nucleotide is unpaired and its structural equivalent (ApoPrdUnPrd).

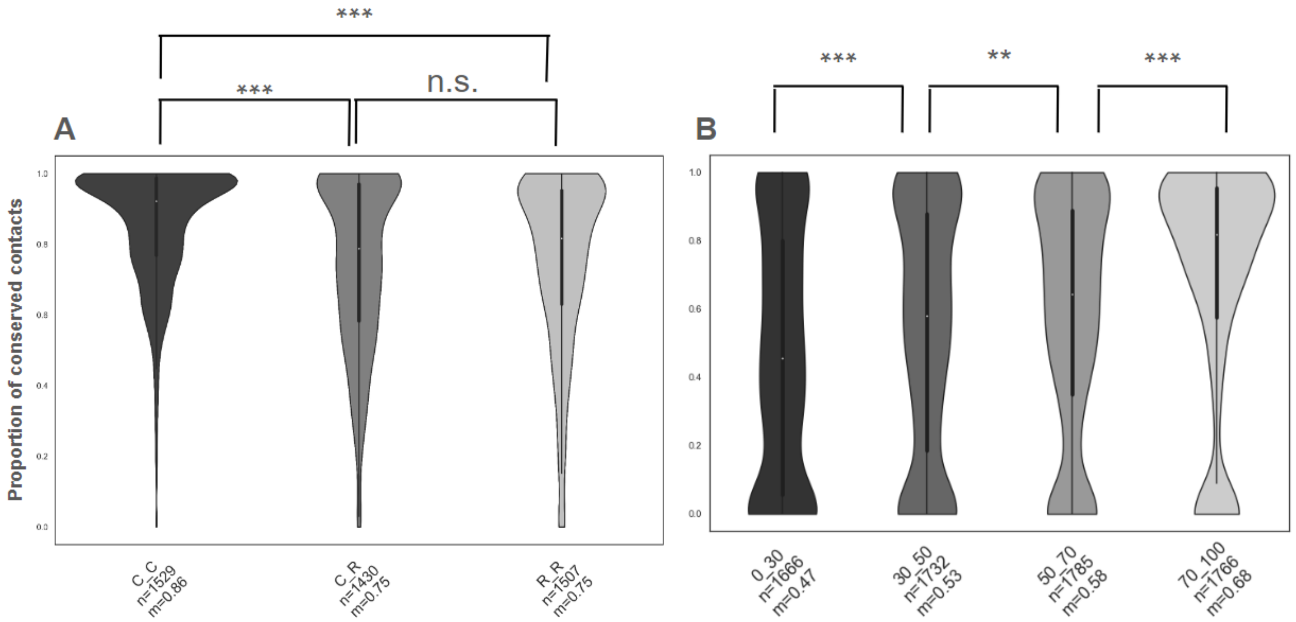


**S5 Fig:** Violin plots of atomic contact conservation depending on amino acid properties. n is the number of interolog pairs used in each violin plot and m is the mean conservation ratio. **(A)** Violin plot distribution of contact conservation for contacts where amino acids belong to the core of the interface in both interologs (C_C), to core in one interolog and rim in the other interolog (C_R) and to the rim region in both interologs (R_R). The differences between C_C and C_R or R_R are statistically significant (p-value < 4e-46 in Wilcoxon rank sum tests) and the difference between C_R and R_R has p-value 0.004 in a Wilcoxon rank sum test. **(B)** Violin plot distribution of contact conservation. From left to right with for the groups [0:30], ]30:50], ]50:70], ]70:100] respectively according to the minimum value of the score within an amino-acid and its structural equivalent. The differences between groups [0:30] and ]30:50], and between groups ]50:70], ]70:100] of conservation scores are statistically significant (p-value<1.6e-6 in Wilcoxon rank sum tests). Between ]30:50], and ]50:70], the difference is less significant (p-value=0.0007 in a Wilcoxon rank sum test).

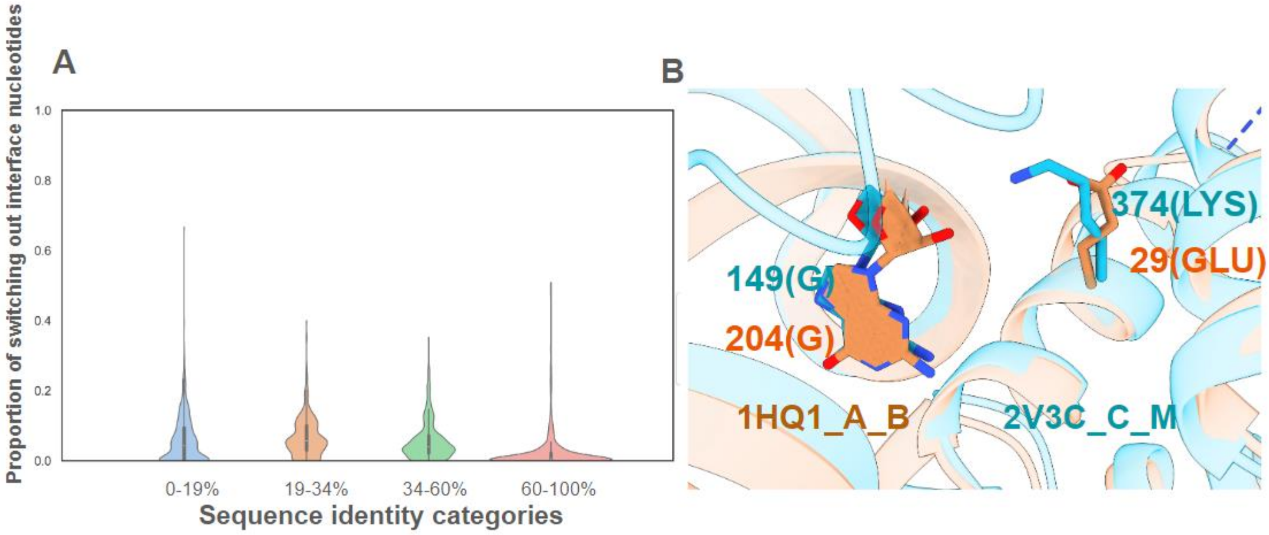


**S6 Fig:** Supplementary analysis of switching out in contact non-conservation. **(A)** Violin plot distribution of the percentage of switching out nucleotides (weighted by the number of atomic contacts in which each nucleotide is involved) across the four ranges of interface sequence identity. **(B)** Illustration of a residue-nucleotide pair [374(LYS): 149(G)] from interface 2V3C_C_M, for which the corresponding pair in the interolog 1HQ1_A_B is no longer a contact, due to amino acid 29(GLU) switching out of the interface.
